# Targeting p16^INK4a^ reverses alveolar epithelial cell dysfunction and induces lung regeneration in emphysema

**DOI:** 10.1101/2025.09.05.674489

**Authors:** Bruno Ribeiro-Baptista, Marylène Toigo, Grégoire Justeau, Perla Abou-Atmeh, Thalles de Freitas Castro, Maéva Zysman, Charlotte Thibaut De Menonville, Etienne Audureau, Klara Schnyder, Huijuan Wang, Yan Hu, Melanie Königshoff, Sophie Lanone, François Chabot, Elodie Zana-Taïeb, Claude Jourdan Le Saux, Mareike Lehmann, Jean-Yves Thuret, Geneviève Derumeaux, Jorge Boczkowski, Janine Gote-Schniering, Laurent Boyer

## Abstract

Pulmonary emphysema involves impaired regenerative capacity of alveolar type 2 epithelial cells (AT2), the main progenitor cells in alveoli. However, the mechanisms underlying dysfunctional epithelial repair remain unclear. In a mouse model of elastase-induced emphysema, we observed an accumulation of activated AT2s in the lung, associated with an overexpression of p16^INK4a^ (p16), a cell cycle inhibitor known to influence stem cell fate. Deletion of p16 promoted the transition of AT2 into alveolar type 1 (AT1) cells, resulting in tissue regeneration in both mice and alveolar organoids. Pharmacological targeting of the p16 pathway using senolytic agents recapitulate this regenerative effect, further supporting the role of p16 as a key brake on epithelial plasticity. These findings demonstrate that alveolar epithelial cell dysfunction can be reversed by p16 deletion or by eliminating p16^+^ cells, thereby reactivating the AT2-to-AT1 transition and promoting endogenous alveolar regeneration. This work identifies the p16 pathway as a promising therapeutic target for restoring damaged alveoli in emphysema.

## Introduction

Pulmonary emphysema is a major component of chronic obstructive pulmonary disease (COPD) characterized by progressive destruction of alveoli and is a leading cause of irreversible chronic respiratory failure. Because no current therapy can restore the alveolar architecture, identifying the mechanisms that enable regeneration of the lung structure in patients with emphysema is of critical importance.

Damaged tissues are repaired by progenitors that are activated during injury. In the alveolar epithelium, a tissue with relatively low renewal capacity, alveolar type 2 epithelial cells (AT2) are one of the main alveolar epithelial progenitor cells, which proliferate and differentiate into alveolar type 1 epithelial cells (AT1)^1^. Recent studies have revealed substantial heterogeneity within the alveolar epithelium, identifying distinct subpopulations of AT2 and intermediate transitional states that orchestrate regeneration^2–4^. AT2 stem cell dysfunction, characterized by impaired progression through these transitional states toward mature AT1, can arise from the combined effects of aging, cumulative environmental insults and genetic predisposition. This dysfunction initiates maladaptive cascades and contributes to the pathogenesis of several alveolar diseases, most notably pulmonary fibrosis^2,5–7^. However, whether similar defects in transitional epithelial populations underlie the impaired regenerative response in emphysema remains poorly understood.

A subset of these transitional epithelial states, particularly Krt8^+^ alveolar differentiation intermediates (ADI), exhibit an upregulation of senescence-associated genes including *Cdkn2a,* which encodes the cell cycle inhibitor p16^INK4a^ (p16). In emphysema, prior evidence has implicated AT2 in an apoptosis - proliferation imbalance skewed toward apoptosis ^8^, along with increased expression of senescent markers ^9,10^. We previously demonstrated in a neonatal disease model that deletion of p16 or ablation of p16 positive cells promoted the recruitment of lipofibroblasts within the AT2 stem cell niche and enhanced alveolar regeneration from early life into adulthood^11^. However, whether targeting p16 in alveolar epithelial cells can similarly restore the regenerative capacity in adult-onset diseases such as emphysema remains unknown.

Here, we hypothesize that epithelial stem cell dysfunction is a key driver of the persistent alveolar destruction in emphysema and that targeting p16 could restore regeneration in the diseased lung. Using an elastase-induced mouse model of emphysema together with human and mouse-derived alveolar organoids, we found that alveolar epithelial stem cell dysfunction marked by p16 overexpression, and accumulation of activated AT2 cells characterized by elevated expression of *Lcn2* and *Lrg1,* contributed to emphysema pathogenesis by impairing lung regeneration. Genetic deletion of p16 reversed this dysfunction, reduced activated AT2 accumulation, and restored alveolar regeneration *in vivo* and in organoid cultures. Moreover, pharmacological clearance of p16⁺ senescent cells using senolytic agents also restored epithelial function and ameliorated emphysema. Together, our findings identify p16-driven epithelial stem cell dysfunction as a central mechanism of emphysema progression and establish targeting p16⁺ cells as a potential regenerative therapeutic strategy in chronic lung disease.

## Results

### Alveolar epithelial stem cell dysfunction in elastase-induced emphysema is characterized by an increase in activated AT2 cells

To assess whether emphysema alters epithelial progenitor capacity, we employed a murine model in which a single intratracheal instillation of elastase produces persistent alveolar injury. This model recapitulated hallmark features of emphysema, including persistent airspace enlargement, as evidenced by a significant and sustained increase in mean linear intercept (MLI) from day 21 through day 150 post-injury (**Fig. 1A–C**).

**Figure 1:**
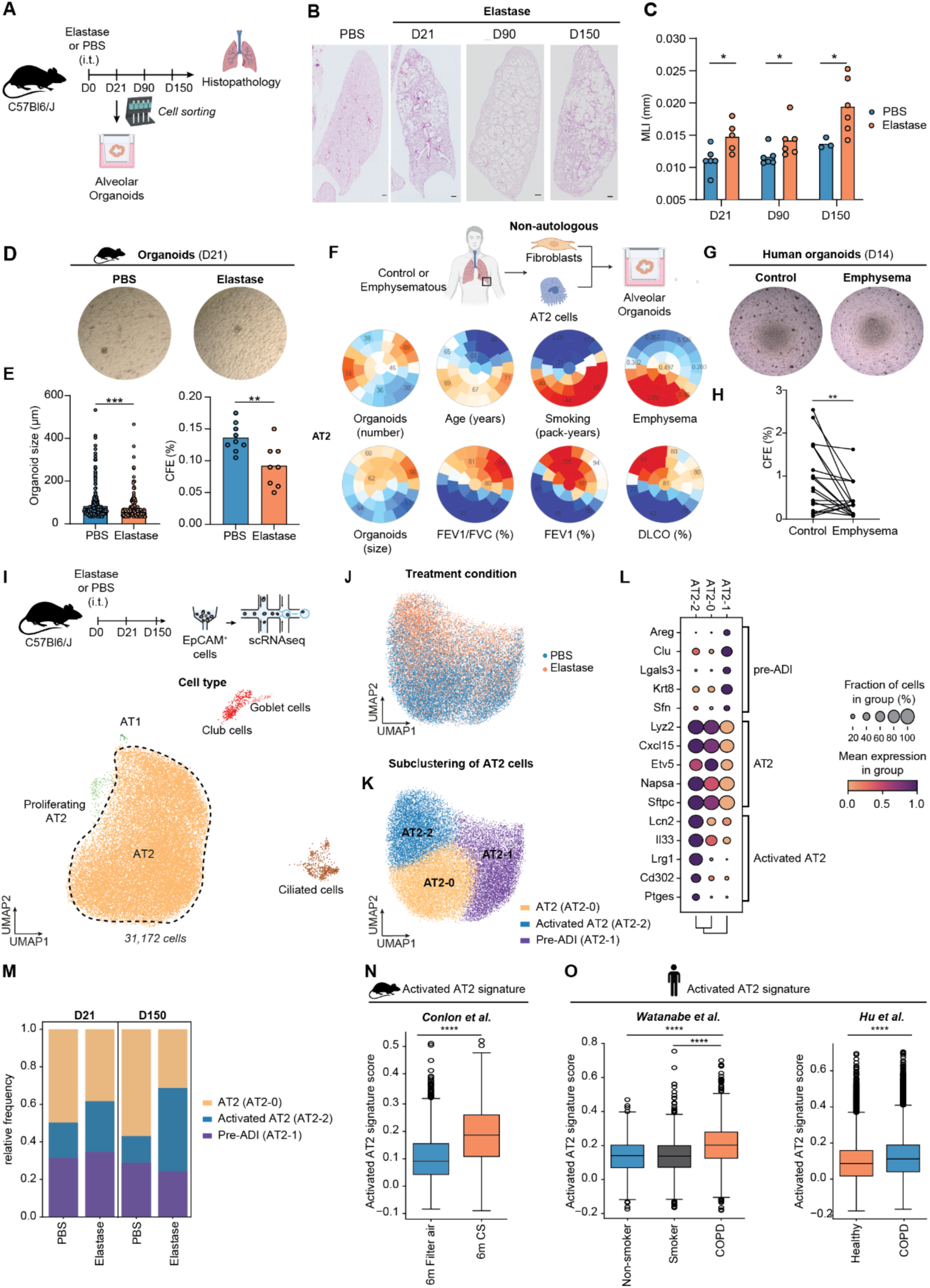
Alveolar epithelial dysfunction and accumulation of activated AT2 in mouse emphysema. (A) Experimental design for lung histopathological analysis and organoid formation assays using isolated lung epithelial cells (EpCAM^+^CD45^-^CD31^-^) from the elastase-induced emphysema mouse model. (B) Representative hematoxylin and eosin stainings of lungs at D21, D90 and D150 post-treatment of elastase or PBS (scale bar = 500 µm). (C) Quantification of mean linear intercept (MLI) measurements (mean is shown, N = 3-6 per group, Mann Whitney test at each time point). (D) Representative images of organoids obtained from co-culture of EpCAM^+^ cells from elastase- or PBS-treated mice with wildtype (WT) fibroblasts after 21 days of culture. (E) Quantification of the colony forming efficiency (CFE, %) and organoid size (µm) (mean is shown, N = 8/group, n > 200 organoids/group, Mann-Whitney test). (F) Exploratory unsupervised analysis of the number and size of formed organoids, patient age, smoking history (in pack-years), proportion of emphysema as quantified by chest computed tomography, FEV₁/FVC ratio (%), FEV₁ (%), and DLCO (%) of human alveolar organoid assays using AT2 from control or emphysematous patients (n=16) and primary fibroblasts from control or emphysematous patients from different donors (n=8). Color intensity reflects the standardized (z-score) value of each variable, allowing visual comparison across patient subgroups, blue represents the lowest average values, and red the highest. Abbreviations: FEV₁/FVC ratio (forced expiratory volume in 1 second / forced vital capacity), FEV₁ (forced expiratory volume in 1 second), FVC (forced vital capacity), DLCO (diffusion capacity of the lung for carbon monoxide). (G) Representative images of human alveolar organoids formed at day 14. (H) CFE as a function of AT2 cell origin. Each data point corresponds to a co-culture of primary fibroblasts with AT2 cells from a patient without emphysema, compared to the culture of the same fibroblasts with AT2 cells derived from emphysematous tissue (N = 18, Wilcoxon matched pairs signed rank test). (I) **S**ingle-cell RNAseq from isolated epithelial cells (EpCAM^+^CD45^-^CD31^-^) of mouse lungs from elastase-induced emphysema model at day 21 and day 150. Uniform manifold approximation and projection (UMAP) showing the cell type annotation. (J) UMAP of AT2 cells clustering depending on PBS or elastase treatment. (K) UMAP of Leiden subclustering of AT2 cells. (L) Dotplot of the marker gene signature of the different AT2 subpopulations. (M) Relative frequency of AT2 subpopulations depending on treatment and timepoint. (N, O) Scoring of the gene signature of the murine activated AT2 cells applied to independent emphysema datasets, including: Conlon et al.^18^ (mouse lungs exposed to six months of filtered air or cigarette smoke, and O Watanabe et al.^19^ (non-smoker, smoker, and COPD patients), and Hu et al.^20^(healthy vs COPD patients). *p = 0.05, **p = 0.01, ***p = 0.001, ****p = 0.0001 and ns = not significant

We first evaluated whether the irreversible structural damage observed *in vivo* reflects a defect in the intrinsic regenerative capacity of epithelial progenitors. To this end, EpCAM⁺ epithelial cells were isolated from control and emphysematous lungs and cultured in organoids with primary lung fibroblasts. Notably, epithelial cells from emphysematous lungs generated significantly fewer and smaller organoids than controls, both at day 14 and 21 of culture (**Fig. 1D–E, Supplementary Fig. 1A**), indicating that emphysema is associated with a durable reduction in epithelial regenerative capacity.

To determine whether these alterations are conserved in humans, we isolated primary AT2 cells by magnetic sorting of HT2-280⁺ cells from explanted lungs of emphysematous and non-emphysematous patients (**Fig. 1F, Supplementary Fig. 1B**) and co-cultured them with primary fibroblasts derived from independent donors (both emphysematous and non-emphysematous). To interrogate the potential relationship between organoid formation and clinical features, we performed an exploratory unsupervised (clustering) analysis using the Kohonen’s self-organized map (SOM) methodology^12^, including the following parameters for both AT2 and fibroblasts patients: number and size of the organoids, age, level of smoking (pack-years), CT scan emphysema score^13^ and lung function parameters (FEV1, FEV1/FVC, DLCO). We observed that organoids were fewer and smaller if AT2 came from emphysematous patients compared to controls based on emphysema score and lung function parameters (DLCO, FEV1) (**Fig. 1F**). However, we did not observe an overlap between organoid size and number with the fibroblast characteristics, suggesting that emphysematous AT2 may be the main driver of organoid formation variation (**Supplementary Fig.1C**). This was further substantiated in paired AT2-fibroblast organoid co-cultures, where emphysematous and non-emphysematous derived AT2 cells were co-cultured with the fibroblasts of the same donor (**Fig. 1G-H**). Emphysematous AT2 yielded significantly fewer organoids than their non-emphysematous AT2 counterparts (n = 18 pairs, **Fig. 1G-H**). Collectively, these data demonstrate that AT2 cell dysfunction is a conserved feature of emphysema in both mice and humans.

To uncover the cellular and molecular programs underlying this defective epithelial regeneration, we next performed single-cell RNA sequencing (scRNA-seq) of lung epithelial cells from control and emphysematous mice. FACS-sorted CD31⁻CD45⁻EpCAM⁺ cells were profiled at day 21 and day 150 post-elastase or PBS treatment, corresponding to early and chronic phases of emphysema (**Fig. 1I, Supplementary Fig. 1D-F**). After quality control, we recovered 31,172 high-quality epithelial transcriptomes spanning all major alveolar and airway epithelial subtypes (AT2, proliferating AT2, AT1, club, goblet, and ciliated cells) (**Supplementary Fig. 1G, Supplementary Table 1**).

Within the alveolar compartment, subclustering of Sftpc⁺ cells revealed three transcriptionally distinct AT2 populations: (i) canonical AT2 cells expressing classical markers (*Sftpc, Etv5, Cxcl15*), (ii) a subset of AT2 cells characterized by *Lrg1*, *Lcn2*, and *Il33* expression, and (iii) a subset defined by *Areg*, *Krt8*, and *Lgals3* expression (**Fig. 1J–L, Supplementary Table 2**). These two last AT2 subpopulation exhibited transcriptional similarity to previously described activated/primed AT2^3,4,14^ and Krt8^+^ transitional alveolar differentiation intermediates (ADI)s/regenerative AT2 cells/DATP/PATS respectively^2–4,6,15,16^, two alveolar epithelial cell states that transiently emerge during specific stages of lung regeneration after fibrotic injury. The similarity of these two subpopulations with activated AT2 and ADI was supported by matchSCore analysis with respective cell state marker genes from a bleomycin-induced lung fibrosis scRNA-seq dataset (**Supplementary Fig. 1I**)^14,17^. Notably, in contrast to ADI/PATS/DATP cells described in fibrosis, these cells retained AT2 identity and were also present in control lungs. We therefore designated this subset as pre-ADI to distinguish it from fully transitional populations. Compositional analysis revealed a selective expansion of activated AT2 cells in emphysematous lungs. Their abundance was increased at day 21 and further enriched by day 150, whereas pre-ADI populations remained unchanged (**Fig. 1M, Supplementary Fig. 1H**).

To ensure that this observation was not restricted to the elastase model, we performed independent validation using external datasets. In a publicly available scRNA-seq dataset of mice exposed to cigarette smoke for six months^18^, we also observed a significant enrichment of our activated AT2 cell signature (**Fig. 1N**). Importantly, projection of the mouse gene signature onto two independent human COPD single-cell datasets^18,19^ confirmed a reproducible accumulation of activated AT2 cells in emphysematous lungs (**Fig. 1O)**. Altogether these results suggest that alveolar epithelial dysfunction, characterized by an accumulation of activated AT2 cells is an important feature of emphysema.

### Alveolar epithelial stem cell dysfunction in elastase-induced emphysema is associated with p16 overexpression

Having established that emphysema is characterized by a shift in AT2 cell state composition, we next sought to define the molecular programs within the AT2 lineage that might underlie their impaired regenerative potential. Differential gene and pathway analyses of the total AT2 population (*Sftpc*⁺ cells) from elastase and PBS-treated mice revealed a marked and persistent transcriptional reprogramming at both early (day 21) and chronic (day 150) stages (**Fig. 2A–B**, **Supplementary Fig. 2A-B, Supplementary Tables 3 and 4**). Elastase treatment induced a shift toward inflammatory and immune-regulatory pathways, including an increase of TNF-α, IL-1β and IL-4/IL-13 signaling, and type I and II interferon responses, signaling axes previously shown to modulate AT2 progenitor function and contribute to maladaptive epithelial repair in chronic lung diseases, including COPD^3,21–23^. Consistent with these pathway changes, many of the top elastase-responsive genes corresponded activated AT2 markers, including *Lcn2*, *Lrg1*, *Cxcl17*, *Cd14*, and members of the uPAR family *Ly6a* and *Ly6i*. Notably, transcriptional regulators of activated AT2 and AT2-to-AT1 differentiation, such as *Tfcp2l1* and epithelial β1-integrin, were not prominently modulated^24,25^. In addition, elastase also altered expression of cell-cycle–related genes, including *Ccnd2* and *Cdca7*, with early induction of *Fos* and *Fosb*, suggesting disrupted proliferative control. Consistent with these observations, signatures of cyclin-dependent kinase pathway dysregulation and increased cellular senescence were detected. Because these findings raised the possibility that senescence may contribute to AT2 dysfunction, we directly assessed senescence signatures within the different AT2 lineage populations. Senescence scores were highest in activated AT2 and pre-ADI cells and further increased following elastase treatment at both time points (**Fig. 2C, Supplementary Fig. 2C-D**). In parallel, expressions of cell cycle associated signatures, including E2F targets, G2/M checkpoint regulators, and mitotic spindle genes were broadly reduced across AT2 subsets (**Fig. 2D, Supplementary Fig. 2C/D**), consistent with impaired cell-cycle progression. As p16 (*Cdkn2a*) is a key regulator of stress-induced cell-cycle arrest and cellular senescence, we examined its expression in elastase-injured lungs. Although *Cdkn2a* transcripts were not detected in the scRNA-seq dataset, immunostaining revealed a marked increase in p16⁺ AT2 cells at all examined time points (**Fig. 2E–F**).

**Figure 2:**
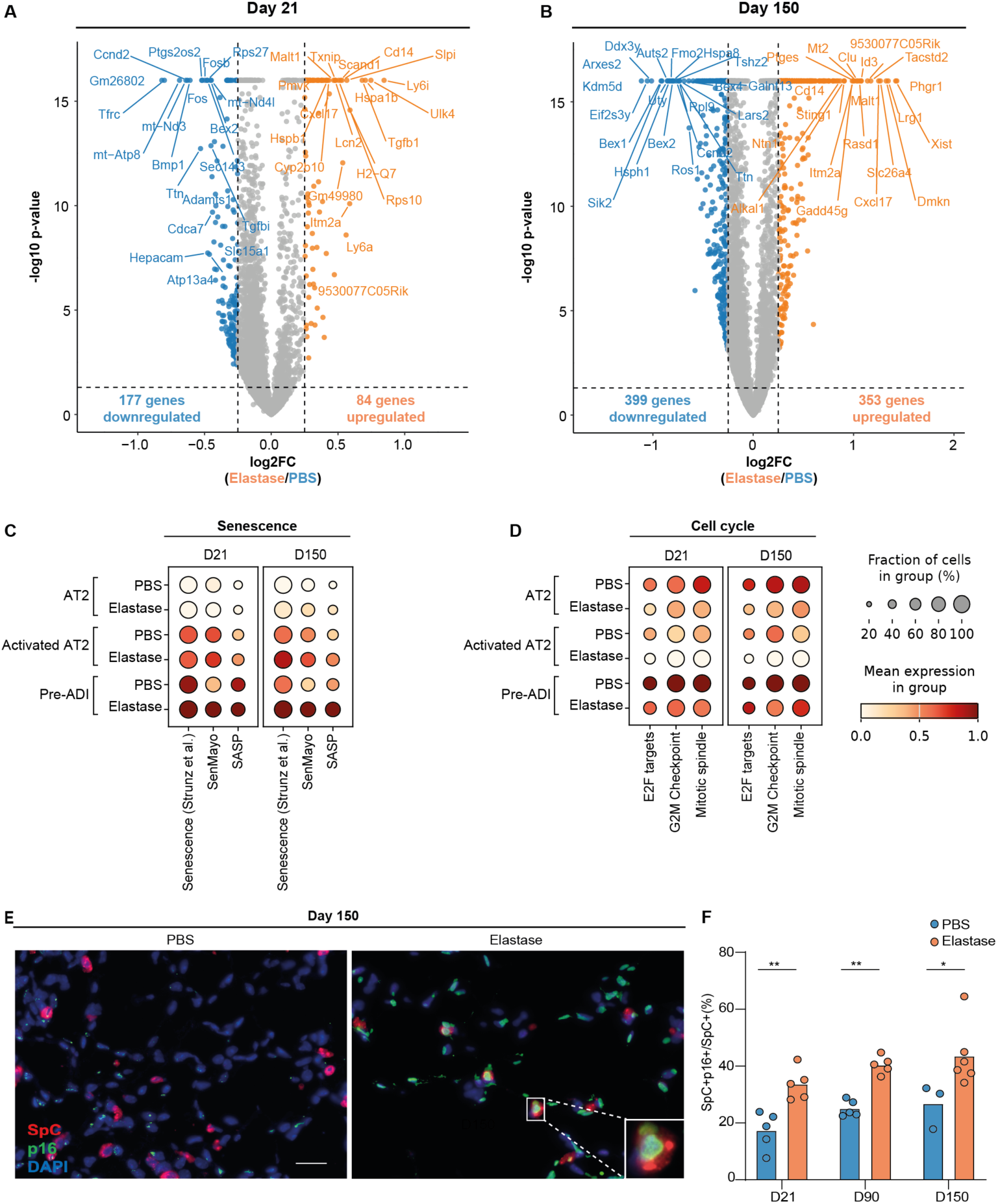
Alveolar epithelial stem cell dysfunction in elastase-induced emphysema is associated with p16 overexpression. Volcano plot of differentially expressed genes at D21 (A) and D150 (B) in AT2 cells of elastase-treated mice compared to PBS controls. A log fold change of 0.25 and q-value of less than 0.05 was used to define differential expressions. Upregulated genes are shown in orange, downregulated genes in blue, and non-significant genes in grey. (C) Dot plot depicting mean scores for three senescence-related gene sets (from^4,26^ and (D) three MSigDB hallmark cell cycle-related gene sets (E2F targets, G2M checkpoint, Mitotic spindle) in the different AT2 subpopulations stratified by treatment condition and timepoint. (E) Representative immunofluorescence staining of pro-SpC (red) and p16 (green) in lungs from PBS and elastase-treated mice. (F) Quantification of p16⁺ AT2 cells as a proportion of total AT2 cells. Data represent mean values (N = 3–6 mice per group). Statistical significance was assessed by Mann–Whitney test at each time point. *p = 0.05, **p = 0.01, ***p = 0.001, ****p = 0.0001 and ns = not significant

Together, these findings identified p16-associated cell-cycle arrest as a potential barrier to AT2-mediated repair, prompting us to test whether p16 deletion could restore epithelial regenerative capacity in emphysema.

### p16 deletion induces alveolar regeneration and reverses epithelial stem cell dysfunction in emphysema

To assess whether indeed p16 overexpression contributes to impaired epithelial repair, we treated WT and p16^⁻/⁻^ mice with elastase and evaluated lung morphology at day 21 (established emphysema), and at days 90 and 150 (persistent emphysema) post-instillation (**Fig. 3A**). At day 21, airway enlargement as quantified by MLI was comparable between WT and p16⁻^/^⁻ mice following elastase injury, indicating that p16 deficiency does not prevent initial alveolar destruction (**Fig. 3B–C**). However, by day 90, p16⁻^/^⁻ mice showed a significant reduction in MLI, indicating spontaneous improvement of emphysema. This regeneration persisted through day 150 (**Fig. 3B–C**).

**Figure 3:**
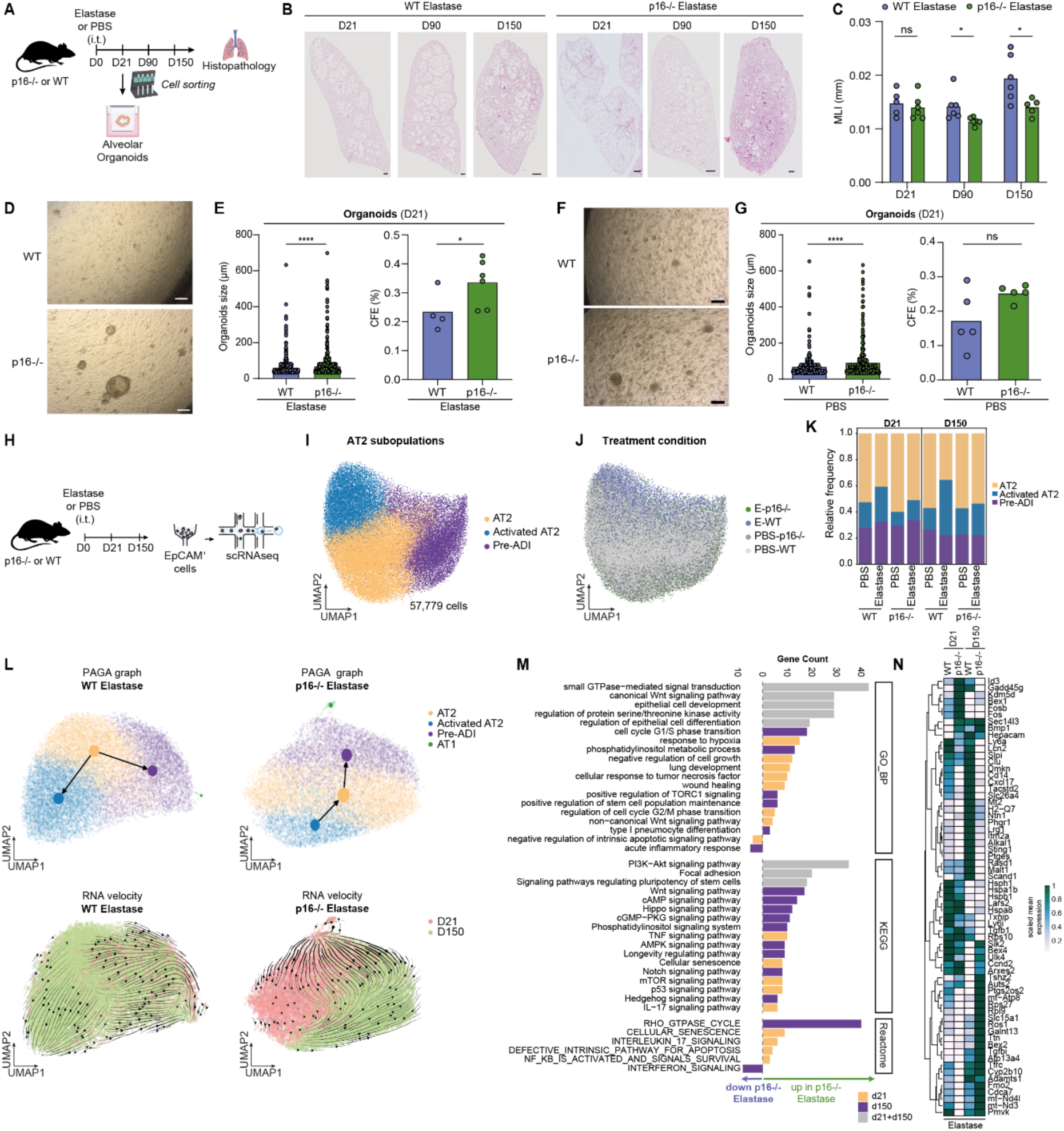
p16 deletion induces lung regeneration and decrease activated AT2 in elastase treated mice. (A) Experimental design for lung histopathological analysis and organoid formation assays using isolated epithelial cells (EpCAM^+^CD45^-^CD31^-^) from lungs of WT and p16^-/-^ mice treated with elastase or PBS, respectively. (B) Hematoxylin-eosin staining of WT and p16^-/-^ lungs at D21, D90 and D150 post-elastase injury (scale bar = 500 µm). (C) Quantification of mean linear intercept (MLI) (mean is shown, N = 5-6, Mann Whitney test at each time point). (D) Representative images of organoids obtained with EpCAM^+^ from WT and p16^-/-^ treated with elastase after 21 days of culture (scale bar = 20 µm). (E) Size (µm) and colony forming efficiency (CFE %) of organoids obtained from EpCAM^+^ cells from WT or p16^-/-^ mice treated with elastase (mean is shown, N>3/group, n > 200 organoids/group, Mann-Whitney test). (F) Representative images of organoids obtained with EpCAM^+^ from WT and p16^-/-^ treated with PBS after 21 days of culture (scale bar = 20 µm). (G) Size (µm) and colony forming efficiency (CFE %) of organoids obtained from EpCAM^+^ cells from WT or p16^-/-^ mice (mean is shown, N>3/group, n > 200 organoids/group, Mann-Whitney test). (H) Experimental design of scRNA-seq profiling in WT and p16^-/-^ mice. (I-J) UMAP of *Sftpc+* AT2 cells depending on cell type annotation. (K) Relative frequency of AT2 subpopulations depending on genotype, treatment, and analysis timepoint. (L) UMAPs with scVelo and PAGA results from WT or p16-/- mice treated with elastase depending on AT2 subpopulations and AT1 (upper panel) or depending on timepoints (lower panel), respectively. (M) Results of GO and pathway enrichment analysis stratified by time point and pathway category comparing AT2 from WT or p16^-/-^ mice treated with elastase. (N) Heatmap of differential expressed genes in WT or p16^-/-^ mice treated with elastase at D21 and D150. *p = 0.05, **p = 0.01, ***p = 0.001, ****p = 0.0001 and ns = not significant

To assess whether improved morphometric outcomes were accompanied by restored epithelial progenitor function, we generated alveolar organoids using EpCAM⁺ cells isolated from WT or p16^⁻/⁻^ mice 21 days after elastase injury and co-cultured them with WT lung fibroblasts. P16 deletion significantly increased organoid number and size (**Fig. 3D-E**), indicating that p16 deletion restored epithelial stem cell function in emphysema. To test whether p16 deletion influences epithelial progenitor potential independently of injury, EpCAM⁺ cells from PBS-treated WT or p16^⁻/⁻^ mice were co-cultured with WT fibroblasts. While organoid numbers were unchanged, organoid size was increased in cultures with p16⁻^/^⁻ cells (**Fig. 3F-G**), suggesting that p16 also modulates baseline epithelial growth potential.

To assess whether fibroblasts contribute to p16-mediated regeneration, we generated organoids using EpCAM^+^ cells from elastase-instilled mice co-cultured with fibroblasts with or without p16 deletion from elastase-treated lungs. P16⁻^/^⁻ fibroblasts produced only a slight, non-significant increase in organoid formation, although overall organoid numbers were low (**Supplementary Fig. 3A-B)**, and p16 deficiency did not increase the fibroblast proliferative potential (**Supplementary Fig. 3C**). Furthermore, immunohistochemistry at day 90 showed no differences in adipose differentiation-related protein (ADRP) staining across genotypes, excluding a role for lipofibroblasts in the regeneration process (**Supplementary Fig. 3D–E**).

Together, these findings indicate that p16 deletion promotes spontaneous late alveolar regeneration primarily through intrinsic epithelial mechanisms, rather than fibroblast-mediated effects.

### Alveolar regeneration is associated with the restoration of alveolar epithelial cell subpopulations

To identify the epithelial populations responsible for the p16-dependent regenerative response, we first examined cell state changes in elastase-injured WT and p16⁻^/^⁻ lungs. We performed scRNA-seq on EpCAM⁺CD31⁻CD45⁻ epithelial cells from WT and p16⁻^/^⁻ mice at days 21 and 150 after elastase injury (**Fig. 3H**). After quality control, we recovered 59,275 epithelial cells, including 57,779 AT2 cells (**Supplementary Fig. 4A)**. The previously defined AT2 subpopulations, i.e. canonical AT2, activated AT2, and pre-ADI cells, were preserved across genotypes, treatments and timepoints (**Fig.3I-J, Supplementary Fig.4B-C**). Strikingly, p16 deletion markedly reduced the proportion of activated AT2 cells in elastase-injured lungs, restoring their abundance to WT baseline levels at both time points (**Fig. 3K**).

**Figure 4:**
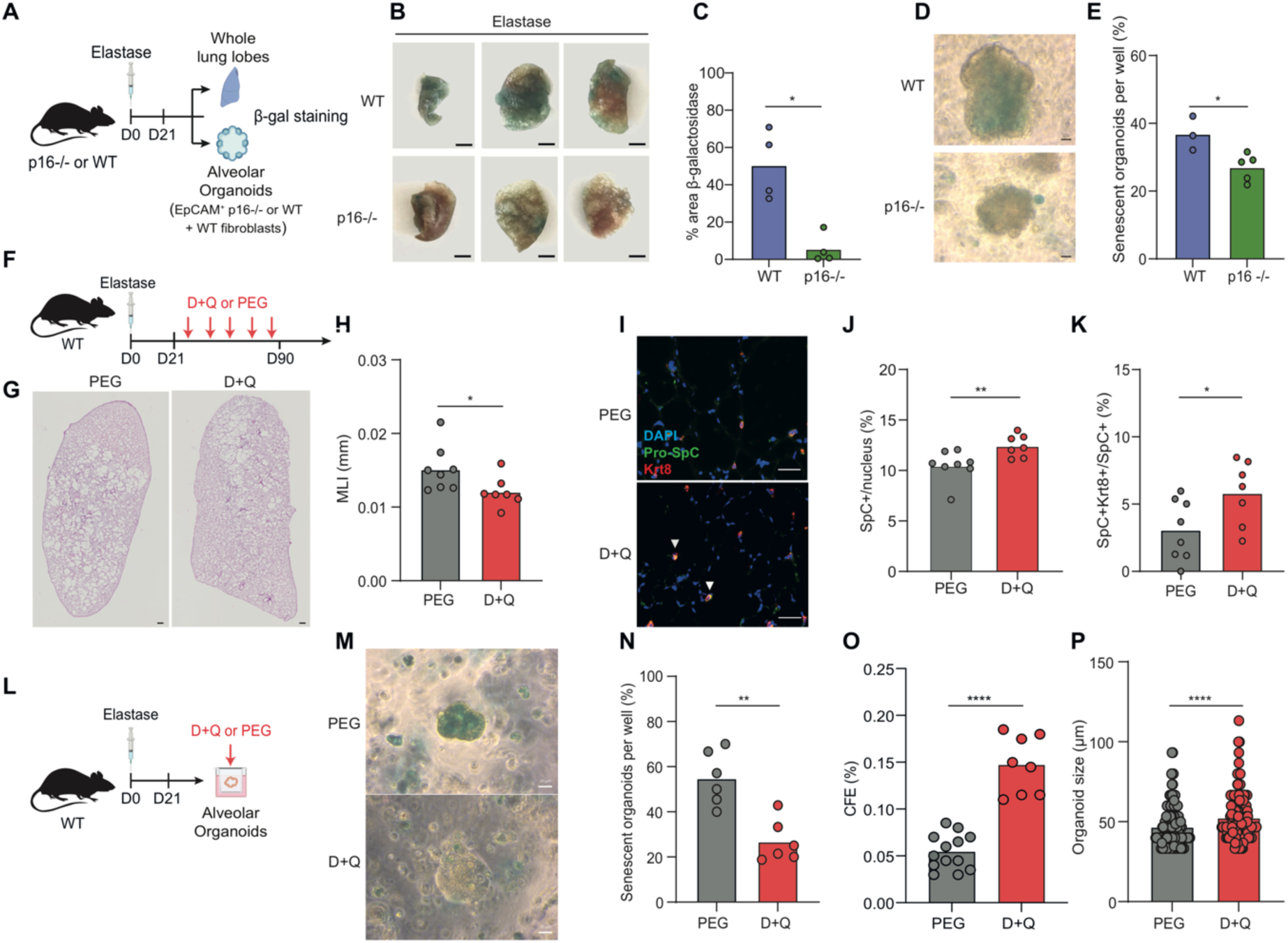
Elimination of senescent cells by senolytics restores epithelial function and reverses emphysema. (A) Experimental design of alveolar organoid formation with EpCAM^+^ cells from WT or p16^-/-^ mice instilled with PBS or elastase cocultured with WT fibroblasts. (B) β-galactosidase activity staining of mouse lungs harvested from p16^-/-^ or WT mice at D21 of elastase instillation (scale bar = 2 mm) and (C) quantification (mean is shown, N = 4/group, Mann-Whitney test). (D) Representative images of β-galactosidase activity staining on organoids (scale bar = 20 µm) and (E) quantification (mean is shown, N >3/group, Mann-Whitney test). (F) Schematic timeline of treatment with senolytics Dasatinib and Quercetin (D+Q) or PEG as control in elastase-treated WT mice. (G) Hematoxylin-eosin staining of the lung at D90 of elastase-treated mice treated with senolytics or PEG as control and (H) quantification of mean linear intercept (MLI) (mean is shown, N=7-8/group, Mann-Whitney test, scale bar = 500 µm). (I) Immunofluorescence staining of Pro-SpC (green) and KRT8 (red) at day 90. (J) Quantification of the number of AT2 cells among all cells (mean is shown, N=7-8/group, Mann-Whitney test). (K) Quantification of the number of Pre-ADI (Pro-SpC^+^/KRT8^+^) cells among AT2 cells (mean is shown, N=7-8/group, Mann-Whitney test). (L) Schematic of organoids treatment with D+Q or PEG from EpCAM^+^ of WT mouse lungs treated with elastase. (M) Representative images of β-galactosidase activity staining on organoids (scale bar = 20 µm) and (N) quantification of the proportion of senescent organoids (mean is shown, N=6/group, Mann-Whitney test). (O) CFE and (P) size of organoids at D14 of culture treated with D+Q or PEG. *p = 0.05, **p = 0.01, ***p = 0.001, ****p = 0.0001 and ns = not significant.

To examine whether p16 deletion restores the AT2-to-AT1 differentiation trajectory, we performed RNA velocity analysis (scVelo) and summarized lineage relationships with partition-based graph abstraction (PAGA). In WT elastase-injured lungs, RNA velocity predicted a differentiation trajectory of AT2 cells toward either activated AT2 or pre-ADI cells in elastase-injured lungs (**Fig. 3L**), indicating a diversion into injury-associated programs with limited progression toward terminal differentiation. By contrast, p16⁻^/^⁻ lungs exhibited trajectories consistent with effective regeneration, with activated AT2 cells progressing toward canonical AT2, pre-ADI, and ultimately AT1 cells (**Fig. 3L**). These computational predictions were supported by immunofluorescence analyses showing increased overall AT2 abundance and expansion of regenerative SpC⁺/KRT8⁺ intermediates in p16⁻^/^⁻ lungs (Supplementary **Fig. 3F-I**).

To elucidate the mechanisms by which p16 deletion reduces activated AT2 cells and restores regenerative capacity, we performed differential gene expression and pathway analyses between elastase-treated AT2 cells from WT and p16^-/-^ animals (**Fig. 3M-N, Supplementary 4D/E, Supplementary Tables 5 and 6**). By day 150, p16 loss resulted in a broad reversal of the transcriptional programs induced by elastase injury. Among the top upregulated genes in elastase-treated WT AT2 cells, including *Lcn2, Lrg1, Cd14, Cxcl17, Id3, Dmkn, Phgr1, Ptges, Clu, Tacstd2*, and *Slc26a4* were strongly downregulated in elastase-treated p16⁻^/^⁻ AT2 cells (**Fig. 3N**). These genes are characteristic of the persistent activated AT2 state, suggesting that p16 is required to sustain this maladaptive program. Conversely, several genes suppressed by elastase in WT mice and associated with AT2 cell cycling, stress tolerance, and differentiation capacity, such as *Ccnd2, Sik2, Auts2, Ttn, Hsph1, Hspa8,* and *Bex4* were upregulated in p16⁻^/^⁻ AT2 cells. Together, these reciprocal changes indicate that p16 deletion shifts AT2 cells out of an arrested, inflammation-driven state and back toward a regenerative trajectory.

Pathway analyses further supported the restoration of AT2-to-AT1 differentiation potential. In p16⁻^/^⁻ AT2 cells, pathways central to epithelial renewal and AT1 maturation, including Wnt and Notch signaling, epithelial cell development, type I pneumocyte differentiation, and stem cell pluripotency programs, were prominently enriched (**Fig. 3M**). Additionally, cell-cycle pathways (e.g., G1/S and G2/M transitions) and pro-survival programs such as PI3K–Akt and mTOR signaling were upregulated, consistent with the increased proliferative and regenerative competence observed in p16-deficient lungs. Notably, enrichment of senescence and p53 signaling pathways was largely confined to day 21, suggesting that p16 deletion reshapes the temporal dynamics of senescence to permit later differentiation and lineage progression.

Altogether, these results suggest that p16 deletion reverses elastase-induced epithelial dysfunction by reducing activated AT2 cells and reactivating programs that restore AT2-to-AT1 differentiation and alveolar regeneration.

### p16 deletion induces alveolar regeneration without aberrant process of fibrosis or cancer

We next questioned if this regenerative process driven by p16 deletion could be in part aberrant with the development of fibrosis or lung cancer^27^. We thus quantified septal thickness (**Supplementary Fig. 5A)**, collagen deposition **(Supplementary Fig. 5B**), and myofibroblasts content **(Supplementary Fig. 5C**) as markers of lung fibrosis. These markers were transiently increased at D90 in p16^-/-^ mice as compared to WT animals and returned to the baseline level at D150, confirming the absence of an aberrant remodeling process and suggesting an important step in the regenerative process involving fibrogenesis. At histological level, we did not observe any pulmonary neoplastic process during the follow-up of mice with a deletion of p16 until D150. Moreover, a CT scan was performed 1 year after elastase instillation on the lungs of WT and p16^-/-^ mice. We did not observe any tumors, also excluding a long-term process of lung carcinogenesis associated with p16 deletion and emphysema (**Supplementary Fig. 5D**).

P53 is a crucial tumor suppressor that suppresses lung adenocarcinoma development from AT2. At the gene level, the emphysematous destruction was associated at D21 with an elevation of p53 and p21 mRNA (**Supplementary Fig. 5E-F**). Altogether, these results show that p16 deletion triggers spontaneous alveolar regeneration in emphysema once lesions are established without the occurrence of fibrosis or lung cancer.

### Elimination of p16+ cells by senolytics restores epithelial function and reverses emphysema

To confirm that deletion of p16 modulates senescence in our mice model of emphysema, we quantified senescence by analyzing senescence-associated βgalactosidase (SA-βgal) staining in lungs from WT and p16^-/-^ mice following elastase injury (**Fig. 4A**). p16^-/-^ lungs showed reduced SA-βgal staining at day 21 (**Fig. 4B, C**). Similarly, organoids derived from EpCAM⁺ cells of p16^-/-^ lungs had fewer SA-βgal–positive structures compared to WT (**Fig. 4D, E**).

To evaluate the therapeutic potential of eliminating p16⁺ cells and reducing senescence, we tested the senolytics Dasatinib and Quercetin in both an *in vivo* emphysema model and organoid assays. WT mice with established emphysema were treated from day 21 to 90 (**Fig. 4F**). Senolytic treatment significantly reduced MLI (**Fig. 4G-H**), increased AT2 cell numbers, and elevated the proportion of pre-ADI cells (SpC⁺/KRT8⁺) compared to vehicle-treated controls (**Fig. 4I–K**). In parallel, we treated with senolytics organoids generated by co-culturing WT fibroblasts with EpCAM⁺ cells from emphysematous WT lungs (**Fig. 4L**). Senolytics reduced SA-βgal staining in organoids (**Fig. 4M-N**) and significantly increased both organoid number and size, indicating improved epithelial progenitor function (**Fig. 4O-P**). Together, these results confirm that targeting p16⁺ senescent cells can restore epithelial function and promote alveolar regeneration in emphysema.

## DISCUSSION

In this study, we show that alveolar stem cell dysfunction in elastase-induced emphysema is associated with an accumulation of activated AT2 cells and p16 overexpression. Genetic deletion of p16 restores epithelial progenitor function and promotes alveolar regeneration. Finally, pharmacological elimination of p16⁺ cells using senolytics reverses both epithelial dysfunction and emphysematous lung destruction. Collectively, these findings identify activated AT2 cells as potential bottlenecks in the regenerative process and establish p16 as a central regulator of their maladaptive behavior, highlighting senescence-targeted interventions as a promising therapeutic avenue for emphysema.

In models of injury-activated lung regeneration, lineage tracing and single-cell based RNA velocity analyses have revealed stage-specific AT2 intermediates that emerge along the AT2-to-AT1 differentiation continuum^2–4^. Early inflammatory primed or activated AT2 populations are followed by ADI/DATP/PATS transitional cells, which are now understood as essential intermediates required for successful alveolar repair. Recent work by Konkimalla et al. demonstrates that ablation of ADI/DATP/PATS cells causes alveolar simplification resembling emphysema, whereas sustained induction of these cells promotes fibrotic remodeling^27,28^. These findings suggest that roadblocks along the differentiation trajectory and imbalances in transitional epithelial states can drive divergent pathological outcomes.

Here, using an elastase-induced emphysema model, we show that activated AT2 cells aberrantly accumulate. Single-cell RNA-seq–based trajectory analyses suggest that in emphysematous lung, the activated AT2 state becomes a dead-end, with minimal evidence of progression toward AT1 differentiation. Strikingly, p16 deletion seems to reopen this stalled trajectory, restoring differentiation toward pre-ADI and AT1 states without inducing fibrosis or neoplastic changes. This aligns with our organoid findings showing improved progenitor function in both injury and homeostasis contexts. These findings suggest that activated AT2 cells in emphysema represent a maladaptive or arrested epithelial state that potentially drives lung pathology.

Hu *et al.* recently discovered a club-cell derived emphysema enriched AT2 subset in both humans (asATII) and mice (ATII-6) with poor alveolar progenitor potential. The cells express markers such as *Ly6i, Lcn2, Slc26a4, Slpi,* genes that were also upregulated in our activated AT2 cell population^20^. We also found evidence of activated AT2 cell signatures in smoking induced emphysema and other human COPD datasets, suggesting a conserved pathological state across mice and humans.

The precise role of activated AT2 cells in driving emphysema progression remains to be determined. Their integration within the alveolar stem cell niche, and their interactions with distal airway–derived epithelial populations, remain poorly understood. It is also unclear whether these cells functionally differ from analogous transitional states seen in fibrosis models or represent an entirely distinct disease-specific phenotype.

Our data suggest mechanistic hypotheses for how p16 enforces the arrest of AT2 differentiation. Canonical regulators of AT2 plasticity leading to activated AT2 signatures, such as YAP/TAZ, Tfcp2l1, or β1-integrin^24,25^, were not significantly modulated in our single-cell analyses. Instead, p16 deletion was for instance associated with a strong induction of GTPase-mediated signaling pathways. GTPases are known to promote epithelial cell proliferation and survival through activation of mitogenic pathways such as MAPK/ERK and PI3K/AKT^29^. These pathways are essential for stem cell expansion and repair following injury. In addition, GTPases integrate extracellular signals, including mechanical signals, to modulate regenerative responses^30–32^, positioning them as key regulators of epithelial renewal that contribute to active maintenance of AT1 cell identity^32^. These findings suggest that p16 programs may intersect with mechanotransduction circuits, thereby imposing a differentiation block on activated AT2 cells.

Another question is whether there are cell-extrinsic signals in the alveolar microenvironment that reinforce the persistence of activated AT2 cells. The work by Konkimalla et al.^27^ highlights the need for precise temporal coordination between epithelial and fibroblast transitional states to rebuild the fibroelastic scaffold during regeneration. In our model, we observed transient accumulation of αSMA+ myofibroblasts at day 90 in p16^-/-^, raising the possibility that disrupted epithelial–mesenchymal balancing may contribute to the persistence of activated AT2 cells or limit access to regenerative trajectories. Future studies are needed to test whether emphysema disrupts these epithelial-mesenchymal crosstalk and how such disruptions feed back onto epithelial fate decisions.

Age further modifies these interactions. We had previously demonstrated that p16 deletion or elimination of p16^+^ cells in the post-natal hyperoxia model of bronchopulmonary dysplasia induced a catch-up alveolar growth between childhood and adulthood^11^. In that model, the restoration of alveolar structure at adulthood was mainly associated with an increase in lipofibroblasts adjacent to AT2 cells after p16 deletion or p16^+^ cells removal. Interestingly, this lipofibroblast expansion was absent in adult emphysema, indicating that the mechanisms by which p16 regulates regeneration differ fundamentally between developmental and adult lungs, even though the regenerative outcome of p16 removal is preserved. The age-dependent divergence between these models underscores the complexity of the role of p16 across developmental stages and suggests that senescence intersects with distinct stromal–epithelial circuits in neonatal versus adult lungs. Nevertheless, in both models, regeneration emerges only at late time points, suggesting that p16 removal resets long-term epithelial trajectories rather than accelerating acute repair.

Finally, our senolytic experiments emphasize the translational potential of these findings. Previous studies have shown that p16 is overexpressed in emphysema, both in mouse models and in human^9,33,34^. Eliminating p16⁺ cells in vivo and in organoids reduces senescence burden, enhances AT2 progenitor activity, increases pre-ADI intermediates, and reverses established alveolar destruction. These results provide compelling evidence that senescence-targeted therapies can restore regenerative capacity even in chronic, advanced emphysema.

In conclusion, our study highlights that targeting p16 in alveolar epithelial cells could be a promising approach to regenerate the lung in emphysema without inducing an aberrant fibrotic of oncogenic processes. Harnessing tissue regeneration in this context could open a novel therapeutic window for patients with severe and irreversible lung disease.

## Methods

### Animals

C57BL/6J mice (8-14 weeks, male and female) were housed in a pathogen-free barrier environment throughout the study (12 h light/dark cycle with access to food and water ad libitum). The Institutional Animal Care and Use Committee approved experimental procedures on mice (APAFIS#25872-2020060317048060v3). The p16^INK4a-/-^ mice were kindly provided by Anton Berns (Netherlands Cancer Institute, Amsterdam)^35^.

### Elastase-induced emphysema

Mice were randomly assigned into PBS or elastase groups. The lung injury was induced by an intra-tracheal instillation of a single dose administration of 2 UI of elastase (Elastin Products Company) diluted in sterile 50 μL of PBS. The control group was treated with sterile PBS only. Mice were sacrificed on day (D)21, D90 and D150 after instillation. All analyses were done blinded.

### Lung morphometry

Murine lungs were perfused with a cannula at 25 cmH2O for 30 min and fixed in 4% paraformaldehyde. Next, they were embedded in paraffin and sectioned at 5 μm thickness. Hematoxylin/eosin staining was performed according to standard procedures. Ten digital photomicrographs at 20X magnification were performed in the histological section. To evaluate the degree of pulmonary emphysema, we used the standard morphometric method of the mean linear intercept as described by Knudsen L et al^36^. Collagen content was evaluated after Sirius red staining as previously described^37^.

### Senescence-associated β-galactosidase staining

Senescence-associated β-galactosidase staining was performed in whole-amount lungs using the Senescence β-Galactosidase Staining Kit (Cell Signaling©). Briefly, lung whole-amount was fixed at room temperature for 45 min in 1 % paraformaldehyde fixative solution, washed three times with PBS, and incubated overnight at 37°C with the Staining Solution containing X-gal in N-N-dimethylformamide (pH 6.0). Whole lungs were subsequently dehydrated with two consecutive steps in 50% and 70% ethanol and embedded in paraffin for serial sectioning. Sections were counterstained with nuclear fast red.

### Immunofluorescence staining

We prepared and stained formalin-fixed, paraffin-embedded 5-μm-thick lung sections using standard procedures. Briefly, sections were deparaffinized through graded Xylene then rehydrated in aqueous solutions with decreasing alcohol concentrations and washed in PBS. Immunofluorescence staining was performed using anti-proSpC (Abcam, ab90716 1/500), anti α-smooth muscle actin (Abcam, ab5694, 1/200), anti-KRT8 (DSBH, TROMA-I, 4 μg / mL), anti-ADRP (Progen GP42, 1/400), anti-p16 (Abcam, ab54210, 1/200). Secondary antibodies (Life Technologies; A11056, A11008, A11001, A11006, A11062, A11012) were incubated for 1 h at room temperature. Nuclei were counter-stained using prolonged anti-fade reagent with DAPI (Life Technologies©, E1740384). Image acquisition was performed using a fluorescence microscope (Axioimager M2; Zeiss) at X40 magnification. For quantitative analyses, randomized images were taken (Axiophoto; Carl Zeiss MicroImaging equipped with a digital camera AxiocamHRc; Carl Zeiss MicroImaging). The quantification of the different stainings was performed using Image J.

### RNA extraction and gene expression analysis

RNA extraction from sorted murine lung cells was done using RNeasy kit (Qiagen©) and quantitative real-time PCR (*q*PCR) analysis was performed using Quant Studio 6 (Applied Biosystems, Illkirch, France) Data were presented as expression relative to hypoxanthine-guanine phosphoribosyl transferase 1 (Hprt 1). For qPCR, the following primers were used : ink4a (fwd: GCAGTGTTGCAGTTTGAACCC; Rev: GTGGCAACTGATTCAGTTGG); Tp53 (Fwd: GGCCCCTGTCATCTTTTGTC; Rev: TGGAGGTGTGGCGCTGA) ; Hprt1 (Fwd: GTTAAGCAGTACAGCCCCAAAATG; Rev: TCAAGGGCATATCCAACAACAAAC).

### Mouse alveolar organoids

Primary lung fibroblasts were isolated from mouse lungs after enzymatic and mechanical dissociation. In brief, lungs were perfused with ice-cold PBS through the right ventricle, excised and minced in small pieces that were placed in dispase (5 UI/ml, Corning), type 1 collagenase (Gibco, 17100-017) and placed in a Gentlemacs Octodissociator (Miltenyi) for 45 min at 37°C. After digestion, the solution was filtered in 70- and 40-micron strainer (Corning), washed and plated in Dulbecco’s Modified Eagle’s Medium/F12 (Invitrogen) supplemented with antibiotic/antimycotic (Thermofischer 15240062) and 10% FBS (Hyclone). Six hours post-plating, supernatant with non-adhesive cells was removed. The purity of culture was assessed by staining for vimentin and the absence of staining for αSMA, CD31, CD45 and cytokeratin. At the 2^nd^ passage, 50,000 fibroblasts were co-cultured with 20,000 EpCAM^+^ cells. These cells were obtained by the same protocol of digestion followed by a negative (Ter119^-^CD31^-^CD45^-^LNGFR^-^CD16/32^-^CD90^-^) than a positive magnetic bead sorting (EPCAM^+^). Cells were resuspended in SABM (small airway basal medium supplemented with vial except epinephrin) and mixed with matrigel (Corning) at a ratio of 1/1 and added to 24-well transwell filter inserts with 0.4 microns pores (Corning) in a 24-well tissue culture plate containing 500 μL of medium (SABM). The medium was supplemented with Rock inhibitor for the 48 first hours then changed every 2 days. Cultures were incubated at 37°C in a humidified incubator (5% CO2). The number of organoids (colony forming efficiency) and the size were analyzed on day 14 and 21 post-plating.

### Human alveolar organoids

Fibroblasts and AT2 cells were obtained from lung tissue from patients undergoing lung surgery for lung cancer from Tenon Hospital, France with documented information and non-opposition for the use of biological waste materials (EP-TN-TU-PLUS-ORG-DE-001). Primary lung fibroblasts were isolated by the explant’s technique as described previously^37^. Briefly, this method involves finely dissecting a small sample of pulmonary parenchyma to remove pleural elements, macroscopic bronchi, and vessels. The dissected tissue is then seeded in a six-well plate with suitable culture media. Over several days, fibroblasts migrate and adhere to the plastic plating. After three passages, all other cell types are eliminated, leaving only pulmonary fibroblasts. AT2s were obtained by magnetic bead sorting using an anti-HT2-280 antibody (Terrace biotech). A suspension of 50,000 fibroblasts and 5,000 AT2 were mixed with matrigel (Corning) at a ratio of 1/1 and added to 24-well transwell filter inserts with 0.4 μm pores (Corning) in a 24-well tissue culture plate containing 500 μL of medium (SABM, Lonza). The medium was supplemented with Rock inhibitor for the first 48 hours then changed every 2 days. Cultures were incubated at 37°C in a humidified incubator (5% CO2). The number of organoids (colony forming efficiency) and the size were analyzed on day 14 post-plating.

### Senolytic drug treatments

Dasatinib (D) and quercetin (Q) were used *in vitro* at 5 μM and 50 μM respectively, and DMSO was used as a control. For the *in vivo* experiment, senolytics were delivered by oral gavage every five days at 5mg/kg for Dasatinib and 50 mg/kg for Quercetin from D21 to D90 after elastase instillation in 10% propylene glycol (vehicle).

### Single-cell RNAseq

For scRNA-seq, a single-cell suspension was prepared from the EpCAM^+^CD31^-^CD45^-^ cells (Anti-CD326: 118213 1/100, Anti-Ter119: 116208, Anti-CD31: 102418 1/100, Anti-CD45: 103155 1/100, DAPI: 422801 1/300, Biolegend) sorted by FACS from the lungs of 2 mice pooled for each group at day 21 and 150 following PBS or elastase instillation in WT or p16^-/-^mice, respectively. Approximately 20,000 cells were loaded onto a 10x Genomics Chip G with Chromium Single Cell 3′ v3.1 gel beads and reagents (3′ GEM v3.1, 10x Genomics). Single index libraries were generated following the manufacturer’s protocol (10x Genomics, CG000204_RevD). The scRNA-seq libraries were pooled and sequenced using a NovaSeq 6000 instrument [read 1: 28 base pairs (bp), Index1 8bp/ Index2 0bp/read2 55bp]. Count matrices were produced using the Cellranger computational pipeline (v7.0.1, STAR v2.7.2a), with reads aligned to the mm10 mouse reference genome (GRCm38). After preprocessing and quality control, scRNA-seq data were analyzed using the Python package Scanpy (> v1.8.2) following the current Single Cell Best Practice Guidelines: https://www.sc-best-practices.org/preamble.html.

### Preprocessing and analysis of epithelial cells scRNA-seq data

To reduce the impact of batch correction on downstream annotations, we first processed and annotated the WT mouse datasets independently, and only afterward integrated them with the p16^-/-^ samples. Quality metrics were calculated using scanpy *sc.pp.calculate_qc_metrics* function. During quality control, genes with fewer than one count or expressed in fewer than ten cells were excluded. Low quality cell filtering was performed using automatic thresholding via the median absolute deviation (MAD) technique, setting 5 MADs for the *log1p_total_counts, log1p_n_genes_by_counts* and *pct_counts_in_top_20_genes* fractions to identify outlier cells. *pct_counts_Mt* were filtered with 5 MADs. Additionally, cells with a percentage of mitochondrial counts exceeding 10 % were filtered out. Doublets were identified using Scrublet^38^ on each sample separately, and ambient RNA contamination was corrected with soupX^39^, with the contamination fraction manually set to 0.4. Normalization was performed using scran’s size factor approach, based on Louvain clustering (resolution = 1.0). Finally, log transformation was applied with the pp.log1p() function in scanpy.

### Highly variable genes (HVGs) calculation and cell type annotations

For each sample, we identified the top 4000 highly variable genes (HVGs) and considered a gene as globally variable only if it was consistently variable across at least four samples. The remaining HVGs were then used as input for Principal Component Analysis (PCA), followed by the construction of a k-nearest neighbor (kNN) graph based on the first 50 principal components. Clustering was performed using the Leiden algorithm, and clusters were annotated based on canonical marker genes. Initially, we carried out a coarse annotation of the dataset and then extracted the EpCAM^+^ epithelial compartment, which included both alveolar and airway cell types, while filtering out contaminant cells from other cell lineages and low quality cells. We then repeated HVG selection, PCA, and clustering to refine the epithelial cell annotation.

### Differential gene expression testing (DGE)

Differential expression analysis was carried out using diffxpy (v0.7.4). A Wald test with default settings was applied to all genes detected in at least five cells. Genes were considered significantly differentially expressed when they showed a log2 fold change greater than 0.25 and an FDR-adjusted *p*-value below 0.05.

### Gene signature scoring

A gene signature for the activated AT2 cell population was derived from the WT AT2 dataset based on a FDR-adjusted *p*-value < 0.05 and a log2 fold change > 2. The final set included the following genes: *Scd1, Lrg1, Lcn2, Itih4, Hdc, Mfsd2a, Ptges, Cxcl17, H2-Q7, 9530077C05Rik, Proz, Phgr1, Alkal1, Slc5a1, Dmkn,* and *Fgg*. In addition, to quantify the expression of senescence and cell cycle pathways, publicly available senescence lists from Strunz et al.^4^, Saul et al.^26^ and Coppe et al.^40^, and cell cycle lists from the MSigDB database were retrieved. Signature scores were computed in Scanpy using the sc.tl.score_genes() function (default settings), by subtracting the mean expression of randomly selected control genes from the mean expression of the signature genes.

### Re-analysis of publicly available single-cell RNAseq data

To assess whether the activated AT2 gene signature was also enriched in other independent scRNA-seq datasets from murine and human lung emphysema, we scored our activated AT2 cell signature on publicly available datasets from COPD patients (GEO GSE173896^19^ and GEO GSE222374^20^), and cigarette smoke exposed mice (GEO GSE151674)^18^. The results were tested for statistical significance using either a Kruskal-Wallis test for multiple or Mann-Whitney U test for two group comparisons. In addition, to assess the transcriptional similarity of the identified AT2 subpopulations with AT2 subpopulations discovered in Bleomycin-induced fibrosis^41^, matchScore analysis was performed as previously described^14^.

### Gene ontology and pathway enrichment analysis

Gene ontology (GO) and pathway enrichment analysis was performed in R using the clusterProfiler package^42^, with libraries from the KEGG, Reactome, and GO Biological Processes and GO Molecular Function databases. Gene sets were considered enriched if the FDR-corrected p-value was below 0.05.

### Trajectory inference analysis

Trajectory inference analysis and fate mapping was conducted using the Python packages scVelo (version 1.10.1) and CellRank (version 2.0.4). We inferred the splicing kinetics of all genes and estimated RNA velocity using the dynamical model in scVelo. We confirmed this finding by inferring the trajectory of AT2 sub-populations using Partition-Based Graph Abstraction (PAGA) analysis.

### Unsupervised analysis

An exploratory unsupervised (clustering) analysis was performed using the Kohonen’s self-organized map (SOM) methodology^12^ to identify if clinical, functional, and radiological characteristics of control and emphysematous patients are linked to the capacity of their AT2 cells or fibroblasts to form organoids. Briefly, this algorithm allows to build 2-dimensional maps from multidimensional datasets, where each map is divided into small groupings (‘districts’) in which data points (here, the co-cultures) are automatically located by the SOM algorithm based on their characteristics: co-cultures with similar features are closely located on the maps, while co-cultures with distinct profiles are farther from each other. Variables considered for this analysis included the number and size of organoids formed, patient age (in years) and gender, smoking history (in pack-years), the proportion of emphysema as quantified by chest computed tomography, the FEV₁/FVC ratio (%), FEV₁ (%), and DLCO (%). The SOMs were generated using the Numero R package^43^ after principal component analysis adapted for mixtures of qualitative and quantitative variables was applied (PCAMix)^44^.

### Statistical analysis

All statistical analyses were conducted using GraphPad Prism (version 10) or the scipy.stats module in Python. The statistical tests used were Mann–Whitney U tests, Wilcoxon matched-pairs signed rank tests, two-way ANOVA, Kruskal–Wallis tests. Data were presented as mean values ± SEM, and results were considered statistically significant if p < 0.05.

## Author contributions

Conceptualization: BRB, MT, GT, MZ, CTDM, SL, JB, JGS, LB

Methodology: BRB, MT, GT, PAA, TDFC, MZ, CTDM, EA, MA, KS, HW, ML, JYT, JB, JGS, LB

Validation: BRB, MT, SL, FC, CJLS, YH, MK, ML, JYT, GD, JB, JGS, LB

Formal analysis: BRB, MT, GT, PAA, TDFC, MZ, EA, CJLS, YH, MK, ML, GD, JB, JGS, LB

Investigation: BRB, MT, GT, PAA, TDFC, MZ, CTDM, EA, MA, KS, HW, ML, JGS, LB

Data curation: BRB, MT, GT, JGS, LB

Supervision: SL, FC, CJLS, ML, JYT, GD, JB, JGS, LB

Funding acquisition: BRB, MT, MZ, FC, GD, JB, LB Writing—original draft: BRB, MT, CJLS, JGS, LB

## Data availability

Single cell RNA-Seq date was submitted to the NCBI GEO database. All code used for data visualization of the scRNA-seq may be requested from the authors upon reasonable request.

## Acknowledgment

This work was supported by a fellowship from INSERM and ARAIRLOR (BRB), UPEC doctoral school (MT), IMRB and Oxyvie (LB), Fondation pour la Recherche Medicale (FRM, LB) and Fondation du Souffle (LB). The authors thank members of Team Derumeaux and Team Lanone for their precious expertise and their help, Fanny Coulpier (Institut Mondor de Recherche Biomédicale, IMRB-INSERM U955) for her assistance in performing the single cell analysis, Denis Mestivier for bioinformatic support (Université Paris Est, Créteil, France), Denis Debrosse (APHP, Service de chirurgie thoracique et vasculaire, Hôpital Tenon, Paris, France) for surgical lung tissues, Slimani Lofti from Université de Paris, URP2496 Pathologies, Imagerie et Biothérapies Orofaciales et Plateforme Imagerie du Vivant (PIV) for CT scan images. The authors also thank Xavier Decrouy, Christelle Micheli (IMRB–INSERM U955, Créteil, France) and Adrien Lalot (animal facility). The assistance of Adeline Henry, Odile Ruckebusch and Aurélie Guguin (Installation de Cytométrie en Flux, IMRB-INSERM U955, Créteil, France). We sincerely thank William J. Zacharias (Cincinnati Children’s Hospital) for his insightful guidance and generous advice regarding organoid generation. Finally, the authors thank Silvia Fre, Bruno Crestani, Philippos Mourikis and for their advice. The Interfaculty Bioinformatics Unit (IBU), University of Bern provided computational infrastructure and support with bioinformatic analyses. We sincerely thank Matthias Brunner for technical assistance.

## Supplementary Figures

**Figure S1:**
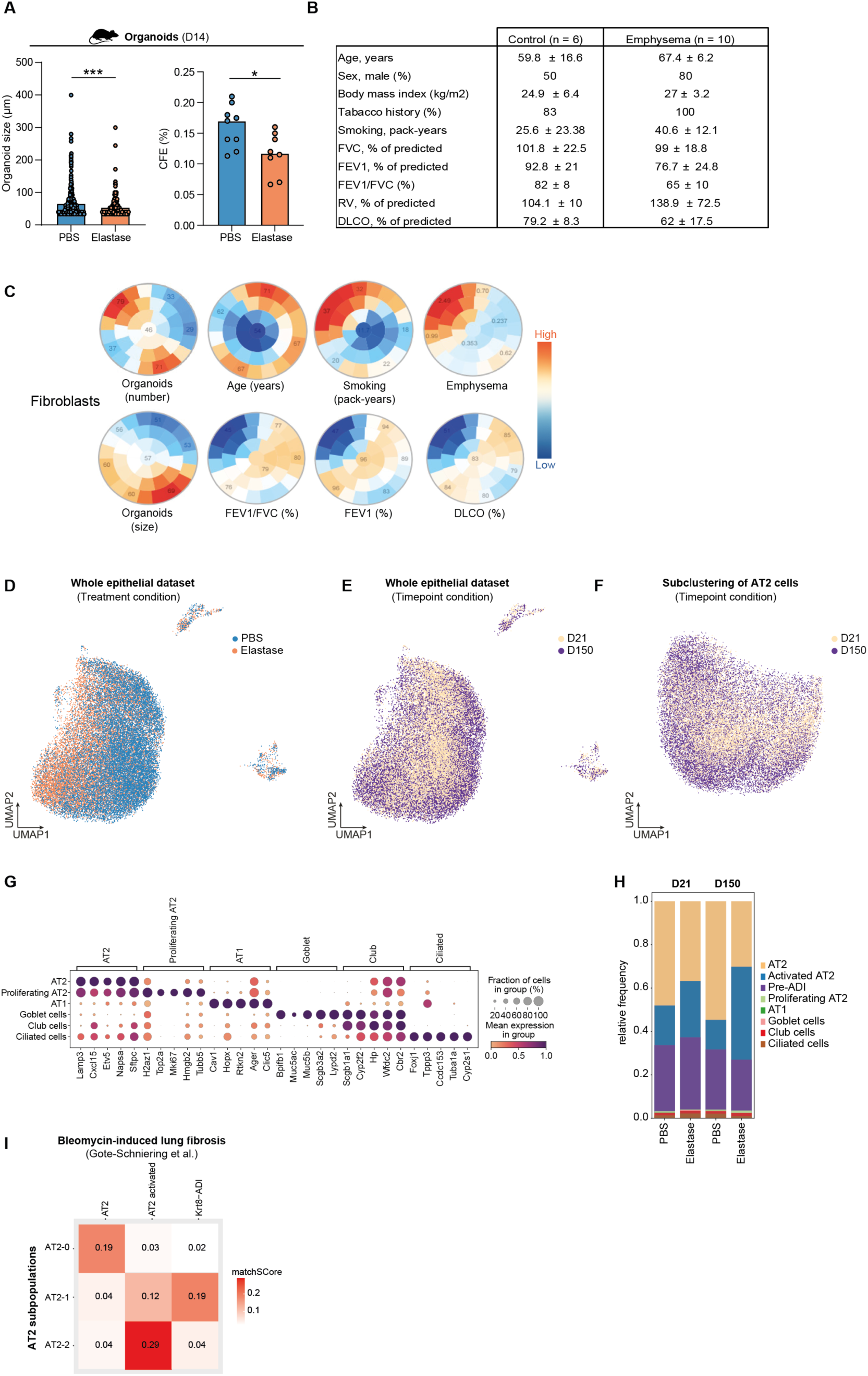
Fibroblasts, epithelial dysfunction, and accumulation of activated AT2 in mouse emphysema. (A) Size (µm) and colony forming efficiency (CFE %) of organoids obtained from the coculture of WT EpCAM^+^ cells obtained by magnetic sorting from elastase and PBS treated mice and WT fibroblasts after 14 days of culture (mean is shown, N = 8/group, n > 200 organoids/group, Mann-Whitney test). (B) Characteristics of patients from whom AT2 cells were used to generate organoids. (C) Exploratory unsupervised analysis of the number and size of organoids formed, patient age, smoking history (in pack-years), proportion of emphysema as quantified by chest computed tomography, FEV₁/FVC ratio (%) and FEV₁ regarding fibroblasts origins. Color intensity reflects the standardized (z-score) value of each variable, allowing visual comparison across patient subgroups. (D-E) UMAP of 31,172 lung epithelial cells from WT mice treated with PBS or elastase depending on treatment and time point condition. (F) UMAP of 29,941 AT2 cells depending on timepoint condition. (G) Dotplot showing marker gene signatures of the entire epithelial dataset. Scaled mean expression is shown. (H) Relative frequencies of cell types/states within the entire epithelium dataset stratified by treatment and time point condition. (I) MatchScore between AT2 subpopulations of our dataset compared with^14^. *p = 0.05, **p = 0.01, ***p = 0.001, ****p = 0.0001 and ns = not significant

**Figure S2:**
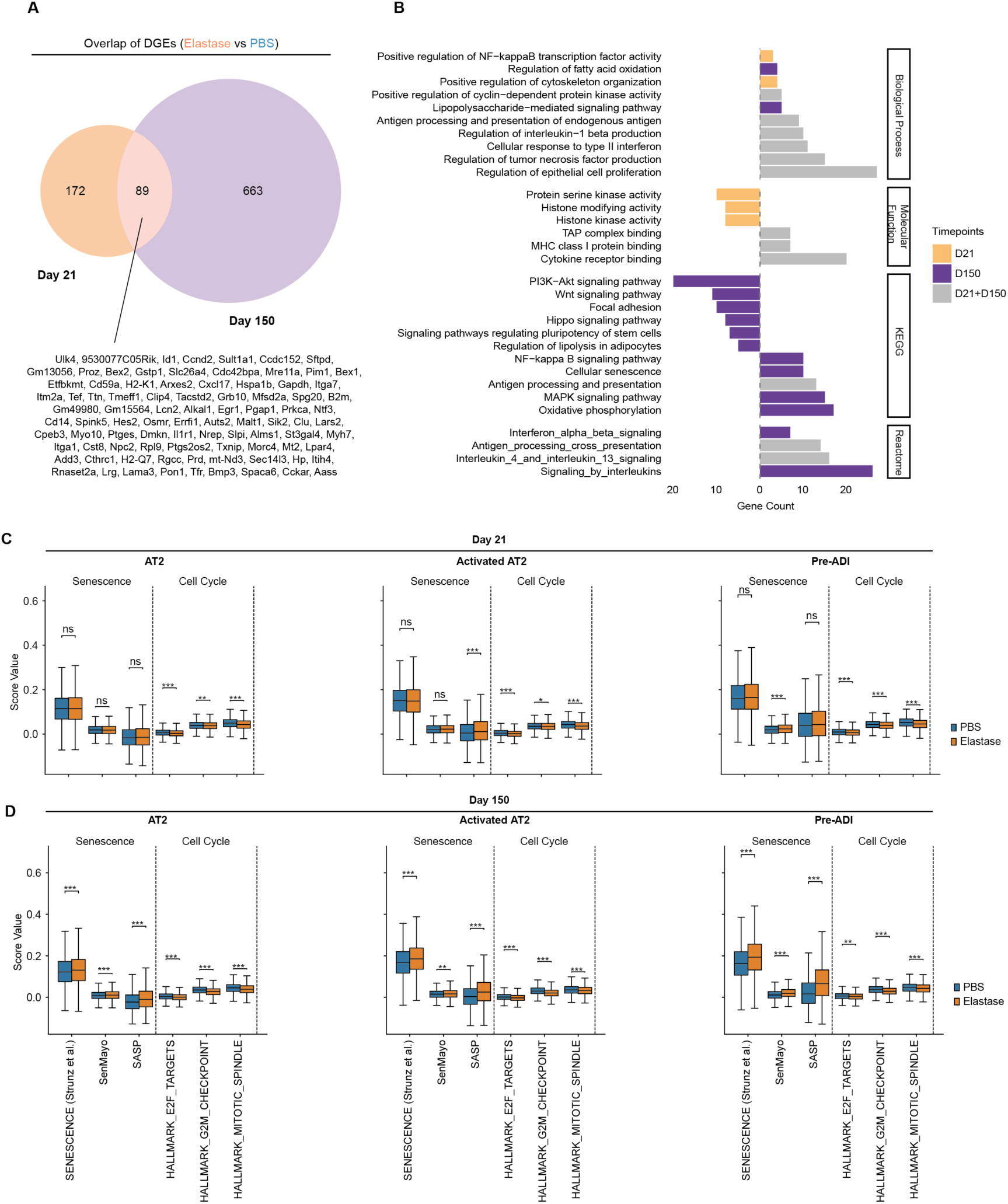
Pathway, senescence, and cell cycle in alveolar epithelial cells in elastase-induced emphysema. (A) Venn diagram showing the overlap of differentially expressed genes (log fold change >0.25, q-value < 0.05) between elastase and PBS-treated AT2 from WT animals at D21 and D150 post-treatment. (B) Representative results of the GO and pathway enrichment analysis stratified by time point and pathway category comparing AT2 from WT mice treated with elastase or PBS. The color code indicates differential regulation at D21, D150 or at both time points (D21+D150). (C-D) Boxplots showing the senescence and cell cycle scoring results across the three AT2 subpopulations at D21 and D150. Statistics: Mann-Withney U test.

**Figure S3:**
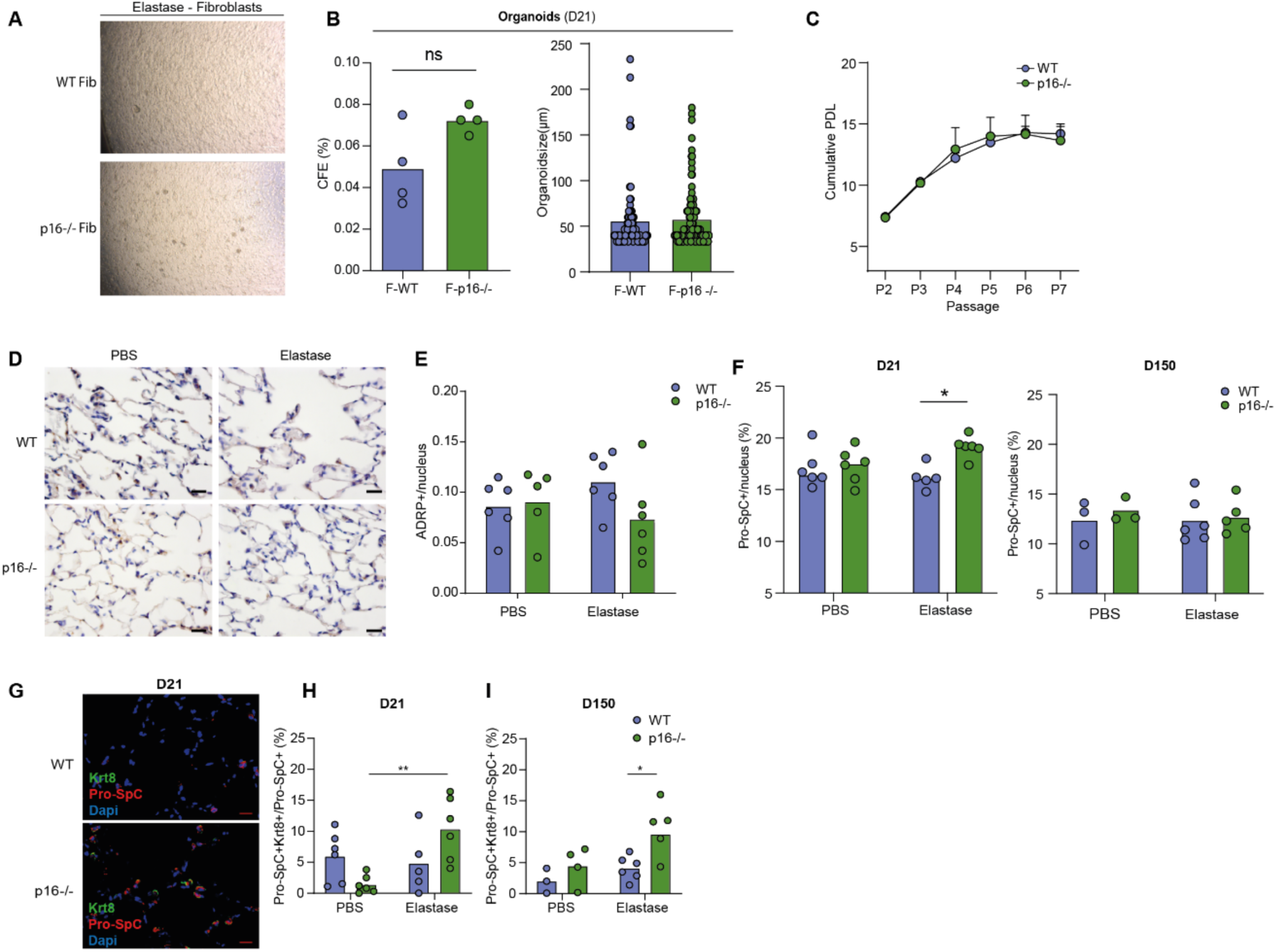
Fibroblasts and pre-ADI in elastase WT and p16^-/-^ mice. (A) Representative images of organoids obtained from the coculture of WT or p16^-/-^ fibroblasts (F-WT and F-p16^-/-^) treated with elastase and WT-EpCAM^+^ cells also treated with elastase (scale bar = 20 µm) (B) Colony forming efficiency (CFE %) and size (µm) of organoids WT or p16^-/-^ treated with elastase (mean is shown, N = 4/group, n > 50 organoids/group, Mann-Whitney test). (C) Cumulative population doubling level (PDL) of fibroblasts from lungs of WT or p16^-/-^ mice from passage 2 (P2) to passage 7 (P7) (n = 3 biological replicates, Error bars indicate the SEM). (D) Immunohistochemistry of lipofibroblasts by adipose differentiation-related protein (ADRP) staining on lungs of WT and p16^-/-^ mice instilled with PBS or elastase (scale bar = 20 µm). (E) Quantification of the number of lipofibroblasts at D90 post-treatment (mean is shown, N = 6, two-way ANOVA test). (F) Quantification of the number of AT2 cells among all cells in WT and p16^-/-^ mice at D21 and D150 (mean is shown, N > 3, two-way ANOVA test). (G) Immunofluorescence of Pro-SpC (red) and KRT8 (green) in WT and p16-/- mice at D21 (scale bar = 20 µm). (H,I) Quantification of the number of Pro-SpC+/KRT8+ among AT2 cells (Pro-SpC+) at D21 (H) and D150 (I) post PBS or elastase treatment in lungs of WT and p16-/- mice (mean is shown, N = 3-6, two-way ANOVA test).

**Figure S4:**
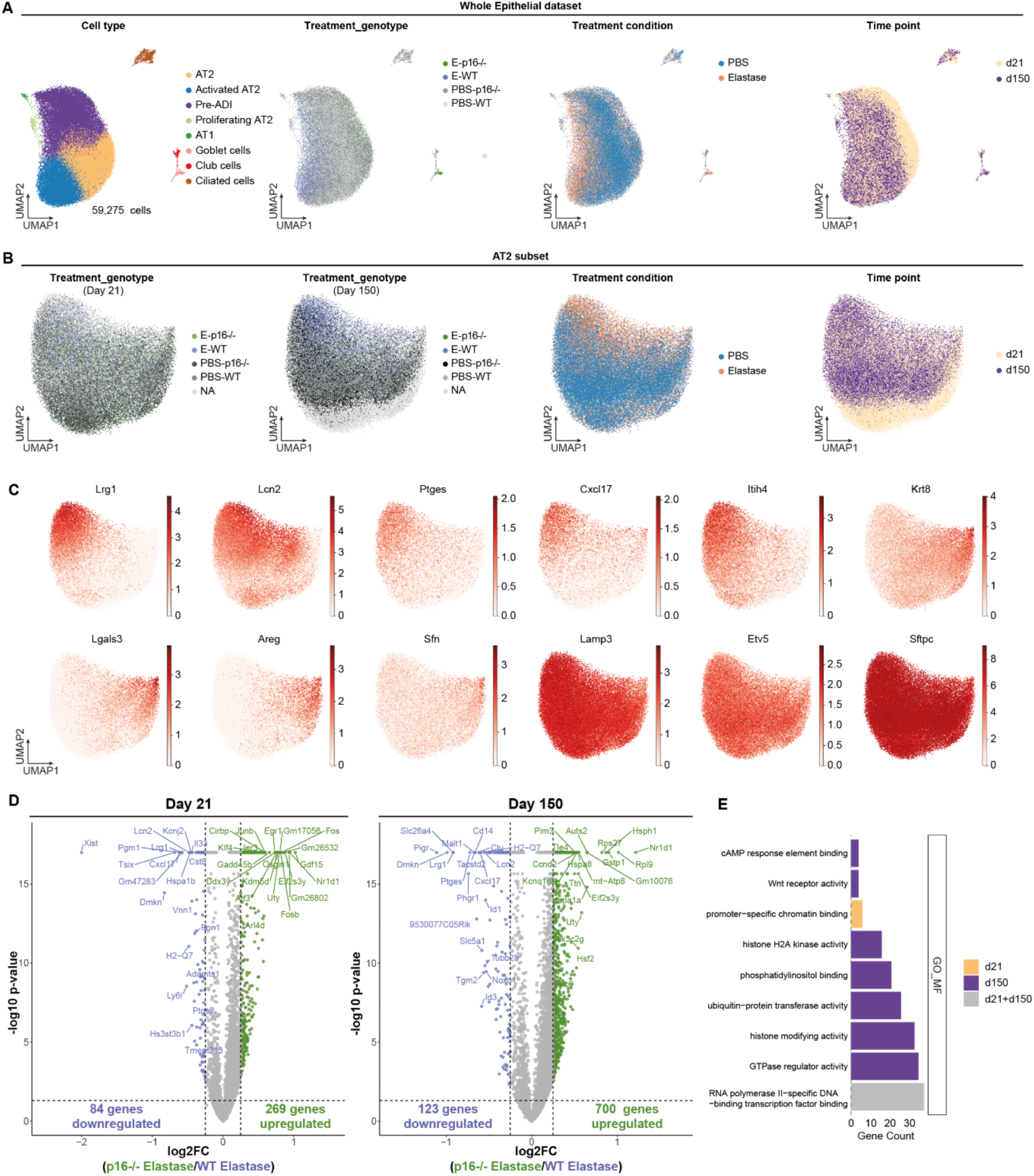
Single-cell RNA-seq of epithelial cells from WT and p16-/- deficient mice treated with PBS or elastase. (A) UMAP of 59,172 lung epithelial cells from WT and p16-/- deficient mice treated with PBS or elastase depending on treatment, genotype and timepoint conditions. (B) UMAP of 57,779 AT2 cells depending on treatment, genotype and timepoint condition. (C) UMAP feature plots highlighting the expression of selected marker genes for activated AT2 (*Lrg1, Lcn2, Ptges, Cxcl17, Itih4*), pre-ADI (*Krt8, Lgals3, Areg, Sfn*), and canonical AT2 (L*amp3, Etv5, Sftpc*). (D) Volcano plots of differentially expressed genes at D21 and D150 in AT2 cells of elastase-treated p16^-/-^ mice compared to elastase-treated WT mice. A log fold change of 0.25 and q-value of less than 0.05 was used to define differential expressions. Upregulated genes in the knockout condition are shown in green, downregulated genes in blue, and non-significant genes in grey. (E) Results of GO Molecular function enrichment analysis stratified by time point and pathway category comparing AT2 cells of elastase-treated p16-/- mice compared to elastase-treated WT mice. The color code indicates differential regulation at D21, D150 or at both time points (D21+D150).

**Figure S5:**
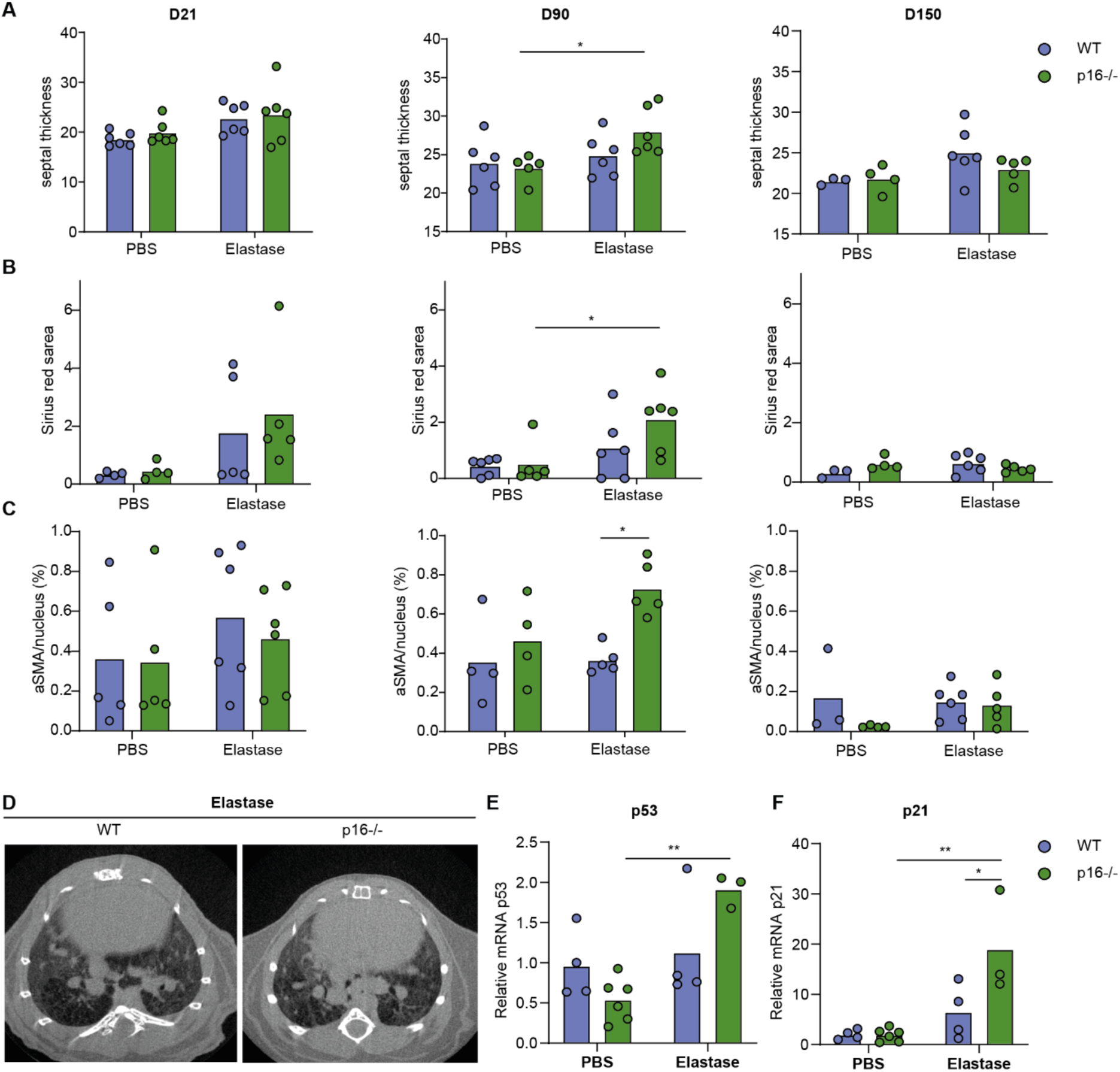
p16 deletion induces lung regeneration without aberrant process of lung fibrosis. (A) Septal thickness of the alveolar wall, (B) Sirius red area and (C) myofibroblasts (αSMA) quantifications at D21, D90 and D150 in WT or p16^-/-^ mice instilled with PBS or elastase (mean is shown, N = 6/group, two-way ANOVA test). (D) Lung CT scan of WT and p16^-/-^ mice 12 months after elastase treatment. (E-F) Relative mRNA quantification of p53 (E) and p21 (F) by RT-qPCR performed 21 days after instillation with PBS or elastase of WT or p16^-/-^ mice (mean is shown, N = 3/group, two-way ANOVA test). *p = 0.05, **p = 0.01, ***p = 0.001, ****p = 0.0001 and ns = not significant

